# Magnetic Resonance Imaging of Gastric Motility in Conscious Rats

**DOI:** 10.1101/2024.09.09.612090

**Authors:** Xiaokai Wang, Fatimah Alkaabi, Ashley Cornett, Minkyu Choi, Ulrich M. Scheven, Madeleine R. Di Natale, John B. Furness, Zhongming Liu

**Author notes:** Correspondence Zhongming Liu, Ph.D. Associate Professor of Biomedical Engineering Associate Professor of Electrical and Computer Engineering College of Engineering, University of Michigan, Ann Arbor.

## Abstract

**Introduction:** Gastrointestinal (GI) magnetic resonance imaging (MRI) can simultaneously capture gastric peristalsis, emptying, and intestinal filling and transit. Performing GI MRI with animals requires anesthesia, which complicates physiology and confounds interpretation and translation from animals to humans. This study aims to enable MRI in conscious rats, and for the first time, characterize GI motor functions in awake versus anesthetized conditions.

**Methods:** We acclimated rats to remain awake, still, and minimally stressed during MRI. We scanned 14 Sprague-Dawley rats in both awake and anesthetized conditions after voluntarily consuming a contrast-enhanced test meal.

**Results:** Awake rats remained physiologically stable during MRI, showed gastric emptying of 23.7±1.4% after 48 minutes, and exhibited strong peristaltic contractions propagating through the antrum with a velocity of 0.72±0.04 mm/s, a relative amplitude of 40.7±2.3%, and a frequency of 5.1±0.1 cycles per minute. In the anesthetized condition, gastric emptying was about half of that in the awake condition, likely due to the effect of anesthesia in halving the amplitudes of peristaltic contractions rather than their frequency (not significantly changed) or velocity. In awake rats, the intestine filled more quickly and propulsive contractions were more occlusive.

**Conclusion:** We demonstrated the effective acquisition and analysis of GI MRI in awake rats. Awake rats show faster gastric emptying, stronger gastric contraction with a faster propagation speed, and more effective intestinal filling and transit, compared to anesthetized rats. Our protocol is expected to benefit future preclinical studies of GI physiology and pathophysiology.

## Introduction

In gastroenterology, evaluating gastrointestinal (GI) motor functions in awake animal models can help translate findings from animals to humans. While clinical trials involving gastric emptying or motility studies are almost always conducted with conscious human participants, most preclinical studies use anesthetized animals. Anesthesia helps stabilize and refrain animals from motion and stress during *in vivo* procedures, such as barostat ^1,2^, serosal recordings ^3,4^, and magnetic resonance imaging (MRI) ^5–7^. However, anesthetics can compromise gastric emptying ^8–11^, myoelectric activity ^12^, intestinal motility and transit ^13–20^. While not fully characterized, the confounding effects of anesthesia are of concern when translating findings from anesthetized animals to conscious humans, especially for high-stake applications, such as testing potential drug therapies.

Among existing methods ^21^, MRI is a promising and non-invasive tool for studying GI physiology and pathophysiology ^22^. It offers rich contrast, balanced spatiotemporal resolution, and a wide field of view, enabling comprehensive assessments of GI motor function in both animals ^5,7,23^ and humans ^24–27^. Our recent work has begun to further unify the acquisition and analysis of GI MRI for rats and humans ^28^. Despite this progress, the need for anesthesia in animals remains a significant hurdle for translational studies. Animal MRI requires body restraints, introduces acoustic noise, and is susceptible to motion artifacts, which pose both technical and physiological challenges when imaging animals without using anesthesia.

A plausible strategy to mitigate anesthesia is to train animals to stay awake and still during MRI, which has been successful for functional MRI (fMRI) of brain activity ^29–41^. The training typically involves acclimating unanesthetized animals to the noise and restraints that occur inside an MRI scanner. However, there has been no prior attempt to use similar strategies for GI MRI in awake animals. Compared to fMRI, GI MRI faces unique challenges due to complex movements of the GI tract alongside respiratory and cardiac motions, which interfere with both image acquisition and analysis of GI motor processes. Awake animals may also experience significant stress due to the acoustic noise and body restraints during MRI, causing unstable physiology ^42,43^. To date, the feasibility of performing GI MRI in awake animals has not been demonstrated, to the best of our knowledge.

In this study, we developed an animal protocol and refined the acquisition and analysis methods for GI MRI in awake rats, extending our prior studies with anesthetized rats ^5,7^. Our goals were to demonstrate the feasibility of GI MRI in unanesthetized rats without excessive body motion or physiological instability, and to establish the awake baseline for gastric emptying and GI motility. By comparison against this baseline, we further assessed the effects of isoflurane and dexmedetomidine.

## Methods and Materials

### Animals and Study Design

In this study, we used 14 Sprague-Dawley rats (SD, male, 230-420 g, Envigo, Indianapolis, IN, USA), with experimental procedures approved by the Unit for Laboratory Animal Medicine and the Institutional Animal Care & Use Committee at the University of Michigan. All animals were housed under controlled conditions with a temperature of 68 to 79 °F, relative humidity of 30% to 70%, and a 12:12 hours dark-light cycle.

Each rat underwent two MRI experiments, first under the awake condition and then under the anesthetized condition (Figure 1A). These two experiments were separated by at least seven days to allow for recovery between sessions. This design enabled us to compare the awake vs. anesthetized conditions for GI motor functions within the same animal. Prior to the first awake MRI experiment, the animals underwent a week-long training period to acclimate them to the MRI environment. During this period, the animals also underwent diet training to condition them to voluntarily consume a labeled test meal for contrast-enhanced GI MRI ^5,7^. Before the second MRI experiment under anesthesia, the animals resumed the diet training but did not undergo further awake acclimation.

**Figure 1:**
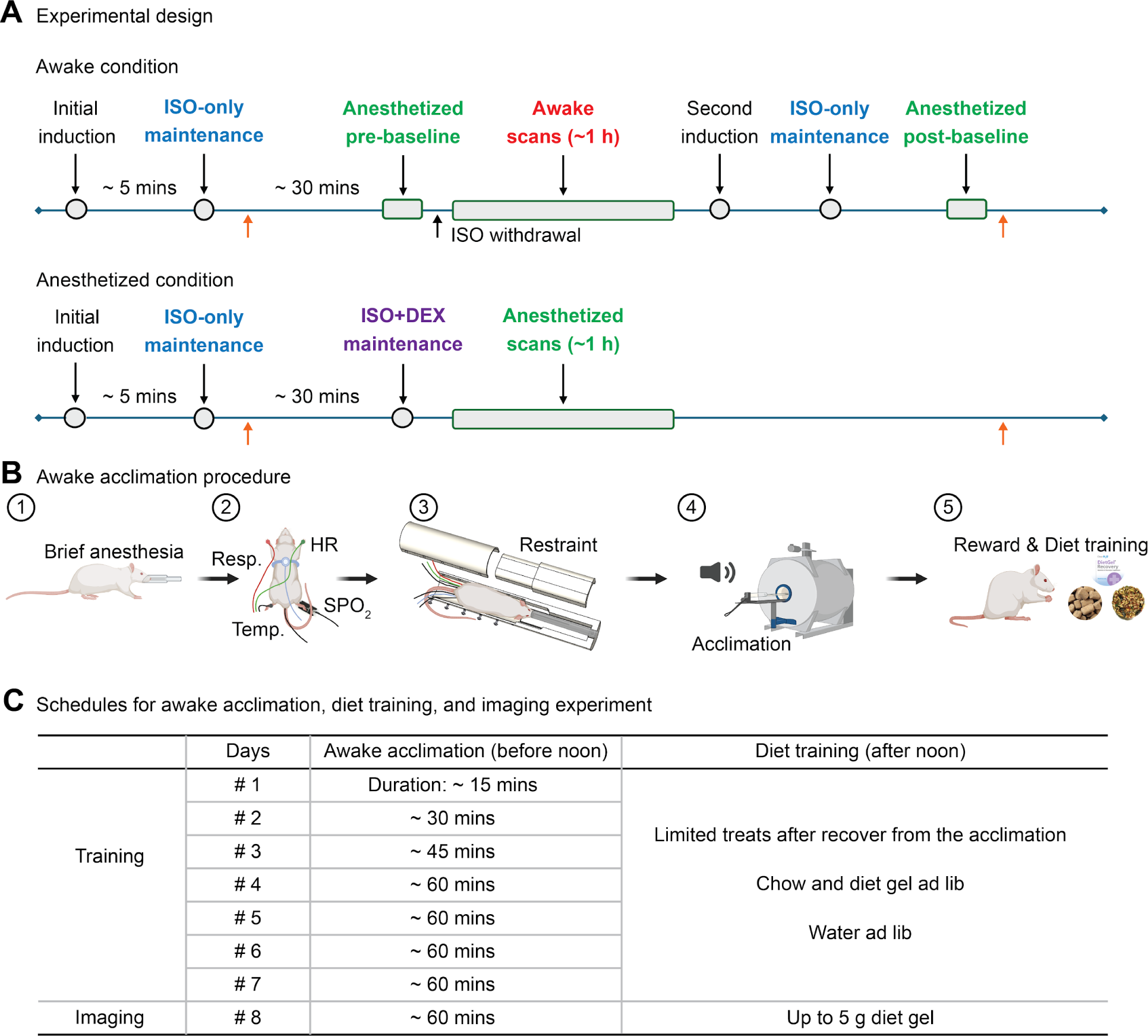
Experimental design and acclimation procedure. **A**. Timelines for the MRI experiments with awake (top) or anesthetized (bottom) rats. Initial anesthesia was performed after the test meal. Orange arrows indicate the time for blood collection. **B**. The procedure for acclimating the rats to the restraints and noise inside the MRI scanner. Physiological recordings of the respiration (Resp), heart rate (HR), temperature (Temp), and pulse oximetry (SPO2), were collected during the acclimation procedure. **C**. Detailed schedules for awake acclimation and diet training over seven consecutive days, and imaging experiments on the eighth day. suu

### Animal Acclimation

We constructed a mock scanner to mimic the real MRI scanner in terms of inner dimensions, acoustic noise, and body and head restraints (Figure 1B). By exposing the animals to this mock scanner for seven consecutive days, we acclimated them to stay awake and still for GI MRI on the eighth day (Figure 1C). Each day around noon, we positioned and restrained the animals in the mock scanner similarly to how they would be set up inside the real scanner. The animals were unanesthetized for increasingly longer periods up to 60 min, further detailed below (Figure 1C). After each session, animals received limited fruit and veggie medley treats as rewards (F7227, Bio-Serv, Flemington, NJ, USA). Following this one-week training, we prepared the animals for the actual GI MRI experiment using the same procedures as those used in the training.

For each training session, a rat was initially anesthetized (up to 5 min) through a nose cone with 5% isoflurane at a flow rate of 1 L min^-1^, and then maintained with 2.5% isoflurane at a flow rate of 0.8-1 L min^-1^. For simplicity, we referred to this anesthesia protocol as “5% isoflurane for induction and 2.5% for maintenance” when describing procedures that used this protocol. We collected blood samples from the tail vein and attached various sensors to monitor vital signs, including respiration, electrocardiogram, pulse oximetry, and body temperature (Model 1030 MR-compatible Monitoring and Gating System for Small Animals, SA II, NY, USA). The respiratory sensor was also used to capture body movements. We then wrapped the animal in soft fabric, restrained it using customized body and head restraints, and placed it into the mock scanner. Pulse oximetry recordings and electrocardiogram were discarded retrospectively because of poor signal quality or incomplete acquisition due to instrumentation errors. The blood collection was only on acclimation days 2, 4, and 6.

We terminated the isoflurane and waited a few minutes until the animal woke up and resumed normal breathing at 80 cycles per minute (cpm) or above. Then, we played acoustic noise (up to ∼90 dB) to simulate the noisy environment during MRI scans. The noise was pre-recorded from the MRI room and included the noise from MR gradient switching, cooling pumps, and ambient sources. After the animal was unanesthetized for up to 60 min, we re-anesthetized the animal with 5% isoflurane for induction and 2.5% isoflurane for maintenance, and removed it from the mock scanner and restraints.

### Diet Training

Through diet training, we trained each animal to voluntarily consume an oral meal (diet gel labeled with Gd-DTPA) for MRI to visualize its content and post-prandial gastric motor responses under a naturalistic feeding condition ^5,7^. The diet training lasted for seven days. Initially, singly housed animals were provided regular chow and diet-gel *ad libitum* (*ad lib*) for the first two days. Subsequently, regular chow was removed from the home cage at 6 p.m. on the second day. From the third to the seventh day, diet-gel was offered at noon with any excess removed at 6 p.m. Water was freely accessible via Lixit throughout the training period.

### MRI Experiments

On the day of an MRI experiment, each animal was fasted overnight for 18 hours, fed a labeled test meal, and scanned for postprandial GI responses while being either unanesthetized (for the first experiment) or anesthetized (for the second experiment). The test meal consisted of 5 g diet gel (DietGel Recovery, ClearH20, ME, USA) mixed with 22.4 mg Gd-DTPA (SKU 381667,

Sigma Aldrich, St. Louis, USA), as described in detail elsewhere ^5,7^. Due to prior diet training, all animals were able to voluntarily consume the test meal. Immediately after the meal, the animals were anesthetized, prepared, restrained, and placed inside the MRI scanner, following the same procedures during awake acclimation.

For the MRI experiment with awake animals, we either terminated the anesthesia immediately after the animals were placed inside the MRI for one group of animals (n=8), or maintained the anesthesia and scanned the pre-awakening GI responses for approximately 10 min before terminating the anesthesia for the other group of animals (n=6). Subsequent procedures were identical for both groups. Shortly after the anesthesia was terminated, every animal increased its breathing rate to 80 cpm (or above), indicating wakefulness. While the animals were awake, we conducted multiple sessions of GI MRI for ∼1 h. Afterward, we re-anesthetized the animals with 5% isoflurane for induction and 2.5% isoflurane for maintenance and performed up to two sessions of GI MRI while they were anesthetized.

For the MRI experiment with anesthetized animals, we did not terminate the anesthesia after placing the animals inside the scanner. Instead, we administered a bolus injection of dexmedetomidine (0.05 mg ml^-1^, Zoetis, NJ, USA) at 0.0125 mg kg^-1^, 0.01 mg ml^-1^, followed by continuous infusion at 0.025 mg (kg*h)^-1^, 0.01 mg ml^-1^ subcutaneously and less than 0.5% isoflurane at a flow rate of 0.8-1 L min^-1^. This combination of anesthesia (referred to as “iso+dex”) was effective in keeping rats stable for MRI ^44–46^. Warm air was blown to the animals to maintain a body temperature of around 37 °C. While the animals were anesthetized (with “iso+dex”), we conducted multiple sessions of GI MRI scans for ∼1 h.

### MRI Protocol

GI MRI was performed using a 7-tesla small-animal MRI system (Varian, Agilent Technologies, California, USA) with a 60 mm volume transmit and receive ^1^H RF coil. We used a multi-slice gradient echo pulse sequence, prescribed 24 oblique slices parallel to the long axis of the stomach, and scanned the entire GI tract every 1.8 s without respiratory gating or <3 s with respiratory gating.

Our acquisition protocol was suited for awake GI MRI because of two technical refinements. First, we used a spatial saturation pulse to suppress surrounding tissues with slow recovery, while highlighting the Gd-labeled intragastric content with fast recovery. This allowed us to achieve rapid acquisition with rate-3 in-plane acceleration while reducing aliasing artifacts. Second, we grouped slices into blocks (2 to 4 slices per block) and acquired all slices in one block before moving to the next. This allowed us to mitigate motion artifacts for each slice acquisition, while using image registration to correct for motion. See more details in the *Motion Correction*.

To scan awake animals without respiratory gating, we used the following parameters: echo time (TE) = 1.8 ms, repetition time (TR) = 6.2 ms, flip angle = 25°, the number of slices = 24, two slices per block, field of view = 64 mm × 42 mm, matrix size = 128 x 84, slice thickness = 1.5 mm. To scan anesthetized animals with respiratory gating, we adjusted the parameters as follows: TE = 1.8 ms, TR = 11.5 ms, and four slices per block.

### Physiological Signal Analysis

We recorded and analyzed animals’ vital signs to assess their physiological state and infer their stress level ^29,44–46^. Each session of the physiological signals lasted approximately 10 min. Briefly, we subtracted the mean and regressed out the linear trend of the signals. We identified the respiratory rate by first band-passing the respiratory recording from 0.25 to 2 Hz and downsampling it to 50 Hz. Then, we detected the inspiration peaks, calculated the inter-peak intervals, and estimated the respiration rate based on the average inter-peak intervals. Similarly, we estimated the heart rate based on the average inter-peak intervals in the pulse oximetry signal filtered from 3 to 10 Hz.

We estimated the heart rate variability (HRV) in terms of the absolute high-frequency power (HF-HRV) ^47^. The inter-peak intervals derived from the pulse oximetry were resampled to 4 Hz with cubic spline interpolation. We further subtracted the mean and regressed out the linear trend from this signal, estimated its power spectral density, and calculated the absolute power from 0.8 to 2.5 Hz ^48^.

Under awake conditions, a stable and non-life threatening physiological state was determined as a respiratory rate of 70 - 115 cpm (or 1.2 - 1.9 Hz) and a heart rate of 300 - 450 beats per minute (bpm, or 5 - 7.5 Hz) ^29,49^. The baseline HF-HRV in awake and free-behaving male rats is in the range of 5.45 ± 3.49 ms^2^ (mean ± standard deviation) ^48^. Under anesthesia, a stable physiological state was determined as a slower respiratory rate of 20 - 60 cpm (or 0.3 - 1 Hz) and a slower heart rate of 250 - 350 bpm (or 4.2 - 5.8 Hz) ^46^. General anesthesia is known to suppress HRV ^50^.

The respiratory sensor could also capture large body movements. When a rat moved inside the body restraint, it pressed the respiratory sensor and corrupted the rhythmicity of the respiratory recording. To quantify this body motion, we calculated the power of the respiratory recording as a function of time and frequency, and identified the time periods with no spectral peak observed in the expected range for the respiratory frequency.

### Image analysis

To analyze GI MRI data, we applied 1) motion correction, 2) image segmentation, and further 3) characterized gastric emptying, GI motility, and transit using both volumetric and surface-based analysis.

### Motion correction

MRI images of the stomach and intestines exhibited movements due to both GI motility and respiratory motion. To correct for respiratory motion, we first denoised the images using a wavelet-based approach ^51^, which reduces noise by transforming the image data into a wavelet domain and filtering out the noise components.

We corrected for respiration-induced global motion first using 2-D affine transformations. Based on the respiratory trace simultaneously recorded with MRI, we identified images collected at the end of expiration and considered them non-motion-corrupted. Specifically, these images were collected when the respiratory signal was within its lowest 10% (end-expiration). These images were averaged to generate the non-motion-corrupted reference. All other motion-corrupted images were then aligned to this reference using 2-D affine transformations.

Next, we isolated residual respiration-induced local motion by averaging affine-corrected images based on the respiratory phases, and modeled the isolated motion by diffeomorphic motion fields. Specifically, we divided each respiratory cycle into 10 phases based on the signal amplitude of the respiratory trace. We labeled each affine-corrected image by the respiratory phase during which it was acquired. By averaging (over time) the images labeled by the same respiratory phase, we obtained a phase series of time-averaged images that only reflected residual respiratory motion, excluding other types of motion due to GI motility.

Using this phase series of images, we optimized a 2-D diffeomorphic motion field to register each respiratory phase to the non-motion-corrupted reference using the Diffeomorphic Demons method ^52,53^. Then we applied the optimized local deformation fields to every image labeled by each respiratory phase. Combined with the optimized affine transformations, these steps co-registered the original time series of images collected at different respiratory phases as if they were all collected at the end-expiration phase, thereby correcting for the respiratory motion (Supplementary Figure 1).

### Gastric and intestinal volume assessment

We evaluated the gastric and intestinal volume changes to characterize gastric emptying and intestinal handling of the food during postprandial processing. To focus this analysis on slow transit rather than rapid motility, we divided the entire series of images into epochs, each including 150 time samples (or 5 to 7 min), and averaged the images within each epoch, to enhance contrast and signal-to-noise ratios. For each epoch, we segmented the time-averaged images using an intensity-based method, followed by manual inspection and refinement, if needed ^7^. From the segmentation, we separately measured the gastric and intestinal volumes for each epoch, and normalized them by the total gastric and intestinal volume of the initial epoch. We resampled such measures by 2 min intervals for each animal and then averaged the results across animals to depict the group-level gastric emptying and intestinal transit. We evaluated the gastric emptying and intestinal transit ratios by their differences within 48 min.

### Surface Modeling and Deformation

We modeled the stomach wall as a surface enclosing the segmented gastric volume, evaluated the dynamic deformation of this surface model, and characterized regional distinctions in contributing to gastric emptying using our recently established method ^7^. Briefly, we applied affine and diffeomorphic transformations to a generic surface template of the rat stomach such that the deformed surface enclosed the gastric content segmented with MRI data in the first or last epoch (or 150 time samples) in the experiment. The difference in the total surface area was used to measure the level of stomach reshaping (shrinkage or expansion) that gave rise to gastric emptying. Since the surface model shared the same topology across times, conditions, and animals, we further characterized the regional deformation, and compared the results between awake and anesthetized conditions ^28^. In particular, we addressed the effects of anesthesia separately on two functional regions: the proximal and distal stomach, as we defined from our previous studies of gastric peristalsis in anesthetized rats ^28^, presumably acting as the pressure (proximal) and peristaltic pumps (distal), which both contribute to gastric emptying ^54^.

### Phasic Contractions of the Stomach

Using the method described in our prior work ^5^, we captured peristaltic gastric contractions from MRI, and characterized their amplitude, frequency, and velocity under both awake and anesthetized conditions. Briefly, we segmented the intragastric volume and identified the axial centerline. We defined cross-sections perpendicular to this centerline and evaluated their area changes over time. We focused our analysis on the distal corpus and the antrum, where the cross-sectional area changes were mostly phasic and periodic, exhibiting a traveling wave propagating along the long axis. Relative to the mean over time, positive area changes reflected muscle relaxations and negative changes reflected muscle contractions. We computed the ratio of the peak-to-peak difference over the time-averaged positive peak, and evaluated its average across time and locations as a normalized measure of contraction amplitude. We measured the contraction frequency based on the maximal spectral power, and measured the propagation velocity by dividing the axial distance between two cross-sections by their difference in the arrival time of the contractile wave.

### Corticosterone Assessment

We collected blood samples on days 2, 4, 6 during acclimation and day 8 (MRI experiments). We centrifuged them at 2,000xG for 10 min, collected serum, and stored the aliquoted samples at −20°C (avoiding freeze-thaw cycles) for corticosterone analysis. Serum corticosterone concentrations were quantified by a competitive corticosterone ELISA (Corticosterone rat/mouse ELISA Cat. No.: RTC002R, Demeditec Diagnostics, Kiel, Germany). We added 10 µLs of neat or diluted serum to each test well for each sample in triplicate. A standard curve (in duplicate) was also tested during each ELISA run. Corticosterone concentrations were measured with a microtiter plate reader (BioTek Agilent Technologies, Inc., Santa Clara, CA, USA) at an absorbance of 405 nm. We discarded blood samples that were less than 50 uL before the corticosterone assessment.

### Statistical Tests

We tested the statistical significance for the difference in gastric motility and emptying between the awake and anesthetized conditions using the paired t-test with a significance level of 0.05. We further evaluated the correlations between different motility features (amplitude, frequency, velocity) and gastric emptying, and tested the statistical significance of each of these factors in driving gastric emptying using ANOVA with a significance level of 0.05.

## Results

We trained 14 rats to stay awake and still inside a 7-T MRI system and scanned them for GI motor responses in either an awake or anesthetized condition, after they voluntarily consumed a contrast-labeled test meal. We evaluated the changes of gastric and intestinal volumes to assess gastric emptying and motility, intestinal filling and transit, and assessed their differences between the awake and anesthetized conditions, as detailed below.

### Stable physiology and intermittent body movements during awake MRI

On the day of MRI, the animals voluntarily consumed 4.1±0.3 g (mean ± standard error of the mean, unless specified otherwise) of the test meal within 21.1±1.7 min. Following the meal, we anesthetized, restrained, and placed the animals inside the MRI scanner. After we terminated the anesthesia, the animals woke up 3.5±0.6 min later, showing more body movements and faster respiration (>80 cpm) (Figure 2). From that time, the animals remained awake for approximately 1 hour, while showing relatively stable physiological signs with a heart rate of 6.0 ±0.1 Hz (Figure 2A), a heart rate variability (HF-HRV) of 8.3±1.2 ms^2^ (Figures 2B), and a respiratory rate of 91.0±1.9 cpm (Figure 2C). These measures are within the normal range for undisturbed and unstressed SD rats ^29,48,49^.

**Figure 2:**
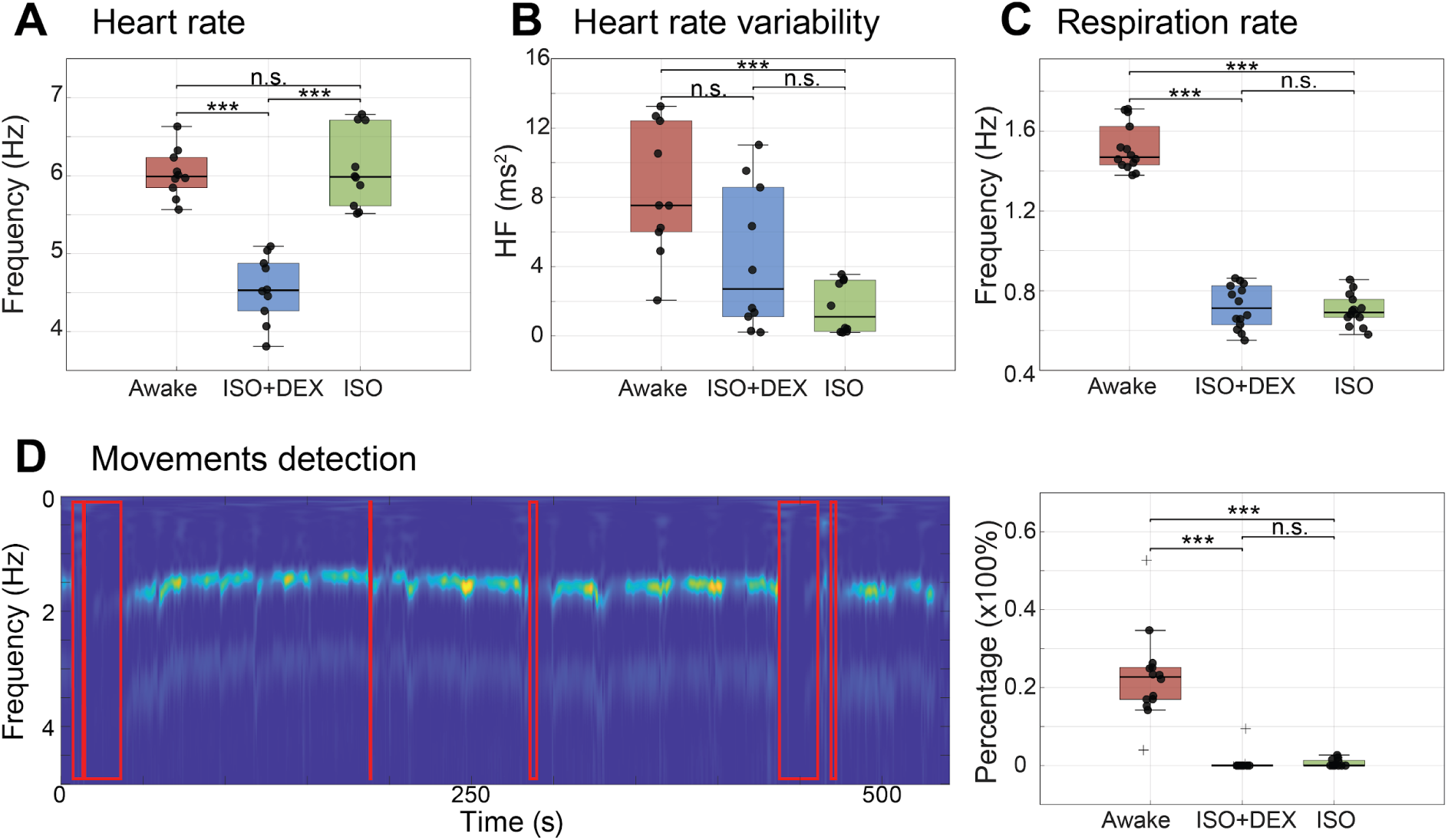
Physiological conditions during MRI in awake and anesthetized rats, in terms of heart rate. (**A**), heart rate variability (**B**), respiratory rate (**C**), and body movement (**D**). Two types of anesthesia were used: ISO+DEX (low-dose <0.5% isoflurane and dexmedetomidine) and ISO (2.5% isoflurane only). In panel **D**, the left shows the time-frequency representation of the respiratory recording with the red boxes highlighting the periods in which body movements corrupted the respiratory signal. The right shows the frequency of occurrence of non-respiratory body movements. The box plot shows the median, 25th, and 75th percentiles, with samples indicated by solid dark circles and outliers indicated by plus signs.

Awake animals exhibited intermittent, but not prolonged, non-respiratory body movements that disrupted respiratory recordings (Figure 2D). On average, the body movement occurred during 21.6±2.8% of the total scan time. As the scan progressed, the movement was less frequent, decreasing from 37.4±4.4% in the first session to 13.2±2.8% in the last session. In contrast, anesthetized rats showed very rare and only small non-respiratory body movements.

Anesthesia affected bodily physiology, depending on the type and dose of the anesthetics used. Compared to the awake condition, 2.5% isoflurane significantly reduced the respiratory rate (42.2±1.3 cpm, p < 0.001) and HRV (1.6±0.5 ms^2^, p < 0.001), but did not significantly affect the heart rate (6.1±0.2 Hz, p = 0.75). The combined use of “iso+dex” significantly reduced the respiratory rate (43.1±1.7 cpm, p < 0.001), the heart rate (4.5±0.1 Hz, p < 0.001), but not HRV (4.4±1.3 ms^2^, p = 0.07).

### Faster gastric emptying and more regional distinction in awake rats

Rats exhibited faster gastric emptying in the awake condition than in the anesthetized condition. Using MRI, we measured the food volume inside the stomach and evaluated its change during digestion, with data available for 48 min in all animals. In the awake condition, the gastric volume decreased by 23.7±1.4%, following a roughly linear trend (Figure 3A & 3E). In the anesthetized condition, gastric emptying was slower, initially linear but almost ceasing after 25 min, giving rise to a 11.2±1.7% volume reduction (Figure 3C & 3F). Key differences between the two conditions are summarized in Table 1.

**Figure 3:**
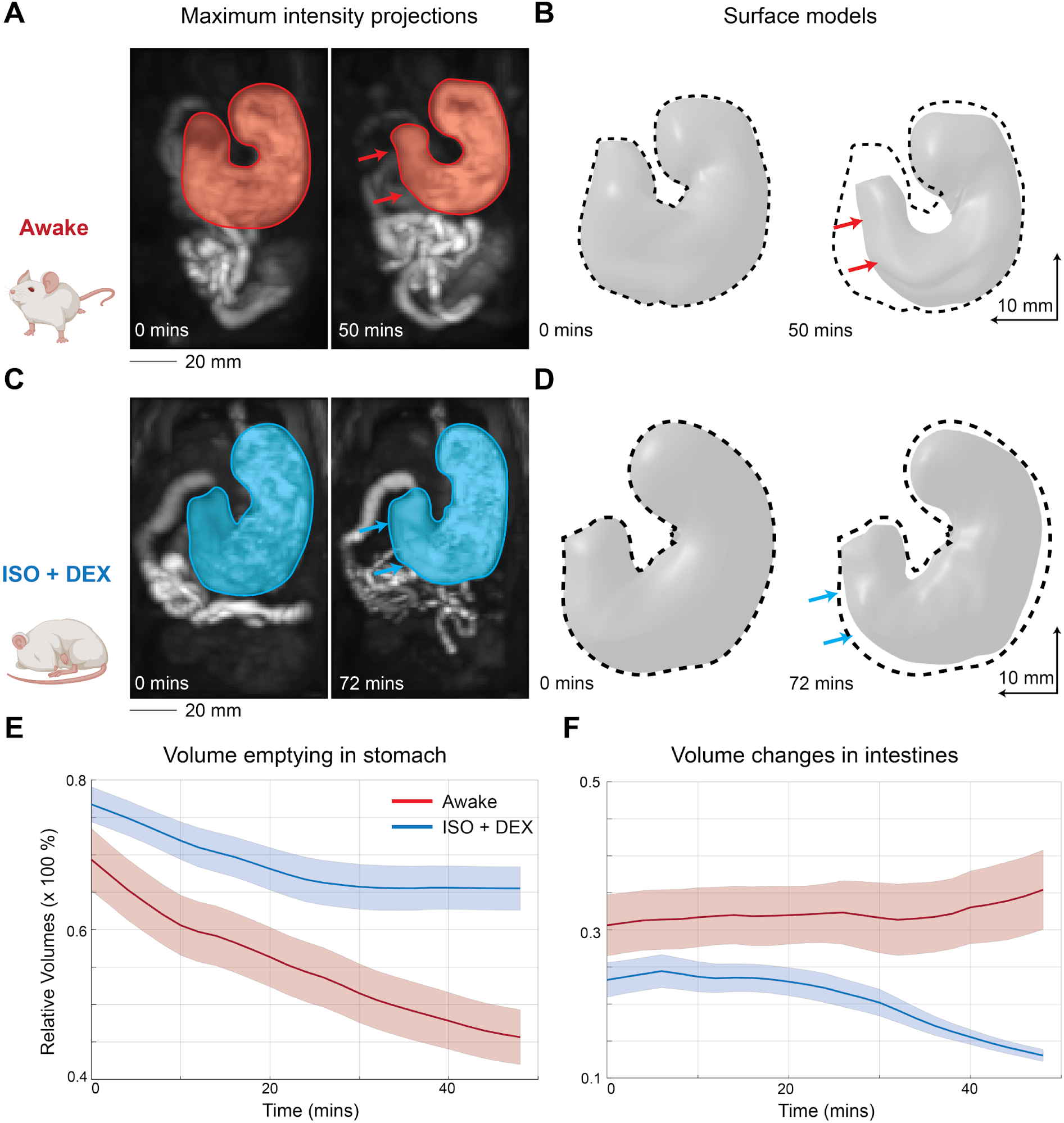
Gastric emptying in awake vs. anesthetized rats. **A** compares the images after maximal intensity projection in the first (0 min) and last (50 min) GI MRI epoch under the awake condition for a single representative rat. Color highlights the segmented stomach. **B** shows the corresponding surface models with the dashed line indicating the contour for the first epoch. **C** & **D** show the results from the same rat in the anesthetized condition (ISO+DEX), using the same format as **A** & **B**. **E** shows the time course of gastric volume changes (in terms of the % of the initial total intraluminal volumes in the stomach and small intestine) averaged across animals, indicating gastric emptying in awake (red) vs. anesthetized (blue) conditions. Similarly, **F** shows the time course of intestinal volume changes averaged across animals.

**Table 1:** Summary of differences in gastric emptying and gastric motility under two conditions. In the group level, the gastric emptying rate was the percentage change of the gastric volume given 48 min after the first scan. Contraction amplitude was the percentage of contraction-induced occlusions. Quantities were expressed as mean ± standard error of the mean. ISO + DEX: low-dose isoflurane and dexmedetomidine.

| | Gastric Emptying Rate (%) | Contraction Amplitude (%) | Contraction Frequency (cpm) | Contraction Velocity ($mm\ s^{-1}$) |
| --- | --- | --- | --- | --- |
| Awake | $23.7 \pm 1.4$ | $40.7 \pm 2.3$ | $5.1 \pm 0.1$ | $0.72 \pm 0.04$ |
| ISO+DEX | $11.2 \pm 1.7$ | $21.7 \pm 1.4$ | $5.4 \pm 0.1$ | $0.56 \pm 0.03$ |

Different regions contributed distinctively to gastric emptying in the awake condition. In a representative rat, we observed regional variation in food retention (Figure 3A) and surface deformation (Figure 3B). The group-averaged surface model confirms this observation with statistical significance (paired t-test, p<0.01) (Figure 4). Given 55 min of gastric emptying, the stomach deformed considerably and reduced its surface area by 466.6±39.2 mm^2^ (or 21.4± 1.7%) (Figure 4D), with seemingly more pronounced changes in the antrum and the proximal fundus (Figure 4B). When dividing the stomach into two functional regions – the proximal and distal stomach (Figure 4A), the proximal stomach reduced its area by 331.8±32.3 mm^2^ (or 15.1 ±1.4%) and the distal stomach by 134.8±15.4 mm^2^ (or 6.3±0.7%, Figure 4E).

**Figure 4:**
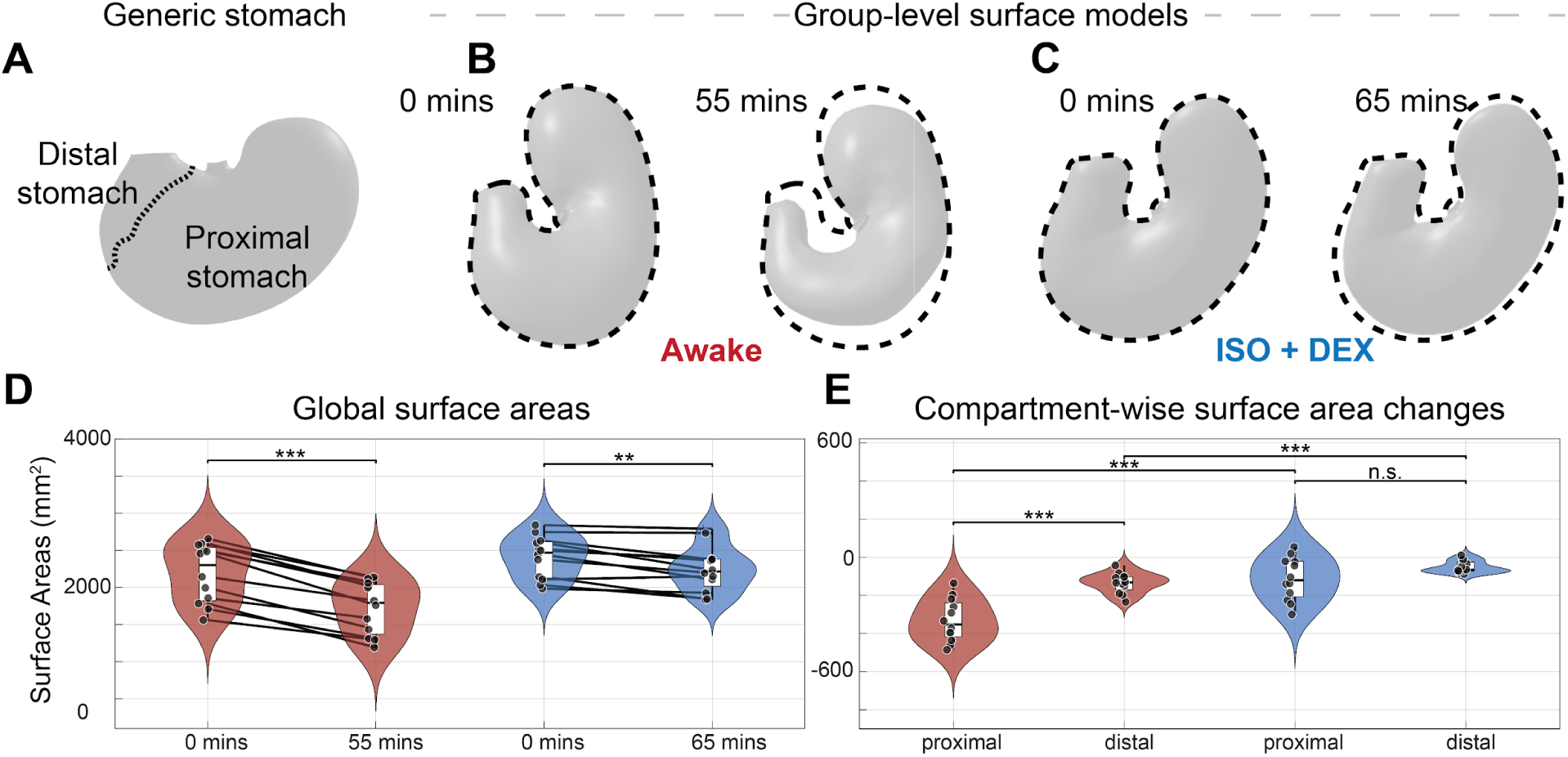
Changes in surface morphology and area in awake vs. anesthetized rats. **A** shows the generic stomach template and its functional division into the proximal and distal stomach by the dashed line. **B** & **C** show the group-averaged stomach shapes at the first and last MRI epoch in the awake & anesthetized condition, respectively. **D** compares the total surface area after about 1 hour for awake (red) and anesthetized (blue) rats. **E** shows the corresponding changes of surface area for the proximal and distal stomachs. The box plots show the median, 25th, and 75th percentiles. The violin plots show the sample distributions. Asterisks indicate statistical significance (paired t-test, * p < 0.05, ** p < 0.01, *** p < 0.001, n.s.: not significant).

Relative to the awake condition, anesthesia affected the stomach’s regional deformation. Given 65 min of emptying in the anesthetized condition, different parts of the stomach appeared to deform uniformly (Figure 4C). As a whole, the stomach reduced its surface area by 164.6±38.1 mm^2^ (6.9±1.6%), approximately one-third of the reduction observed in the awake condition (Figure 4D). The proximal stomach was reduced by 113.0±32.4 mm^2^ (4.7±1.3%) and the distal stomach by 51.6±9.1 mm^2^ (2.2±0.4%), showing less regional distinction compared to the awake condition (Figure 4D).

### Stronger and faster propagation of gastric peristalsis in awake rats

In the awake condition, the rat’s stomach maintained strong contractions throughout the distal stomach. On average, the peristaltic contraction had an amplitude of 40.7±2.3%, a frequency of 5.1±0.1 cpm, and a propagation velocity of 0.72±0.04 mm s^-1^. Dynamic MRI showed that the peristaltic contraction was strong on both the greater and lesser curvatures, often occlusive, effectively moving the content inside the stomach (Figure 5A, B, & C).

**Figure 5:**
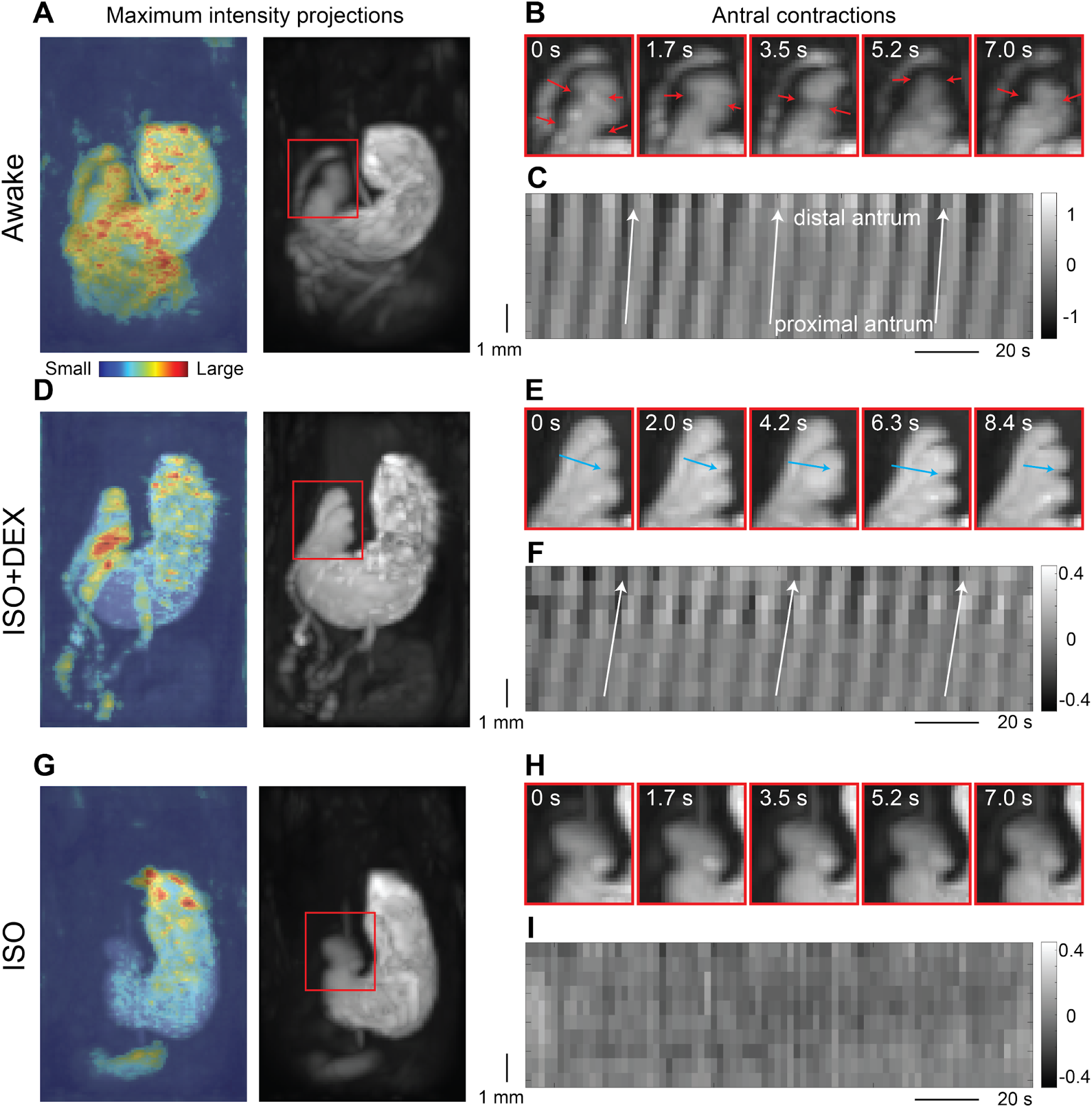
Gastric peristalsis in awake and anesthetized conditions. **A**, **D**, and **G** show the MR images with maximal intensity projection for a representative rat in the awake condition and two types of anesthetized conditions: low-dose isoflurane and dexmedetomidine (ISO+DEX) and isoflurane only (ISO), respectively. The color overlay indicates the temporal variation of image intensity. **B**, **E**, and **H** zoom into the corresponding images of the antrum highlighted by red boxes in **A**, **D**, and **G**, respectively. Arrows in color indicate contractions. Note that panel **B** shows more frame-to-frame variation than panel **E** or panel **H**. **C**, **F**, and **I** show the motility profiles, visualizing the normalized cross-section area changes from the proximal antrum to the distal antrum over time. Only for visualization, each row has its mean removed and divided by its time average. White arrows indicate the propagation of the gastric peristalsis, with its angle indicating the propagation velocity. A more vertical line indicates a faster propagation velocity.

Relative to the awake condition, anesthesia attenuated peristaltic contraction and slowed its propagation. With light anesthesia maintained by “iso+dex”, antral contractions were generally weaker and relatively more evident on the lesser curvature than the greater curvature (Figure 5D, E, & F). On average, the contraction had an amplitude of 21.7±1.4%, a frequency of 5.4±

0.1 cpm, and a velocity of 0.56±0.03 mm s^-1^. Both the amplitude and the velocity were significantly lower than those in the awake condition (paired t-test, p < 0.001), whereas the frequency was similar (p = 0.12). In contrast, the effects of 2.5% isoflurane without dexmedetomidine were even more pronounced, largely disrupting, if not entirely diminishing, peristaltic contractions and food transit within the stomach (Figure 5G, H, & I). See Supplementary videos 1, 2, and 3 for dynamic MRI in a representative rat, demonstrating the different patterns of GI motility in the awake vs. anesthetized conditions. The amplitude of peristaltic contraction appeared to be the major factor contributing to the different rates of gastric emptying across animals and conditions (Figure 6). Gastric emptying was positively and significantly correlated with the amplitude of peristaltic contraction (r = 0.74, p < 0.001) (Figure 6D), but not with its frequency (Figure 6E) or velocity (Figure 6F).

**Figure 6:**
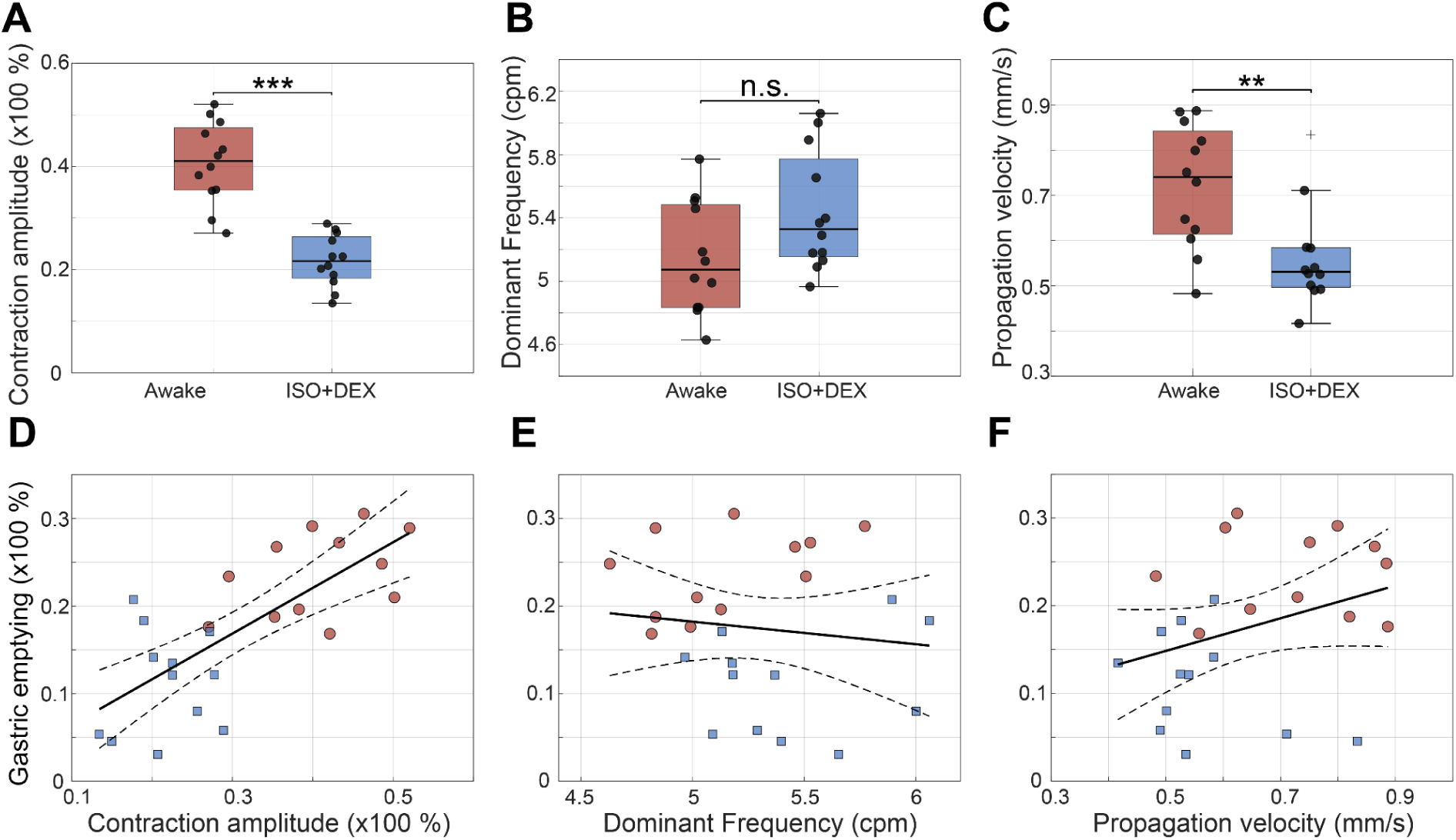
Features of gastric peristalsis and their associations with gastric emptying. **A**, **B**, and **C** show the group-level statistics for the amplitude, frequency, and velocity of gastric peristalsis in the awake vs. anesthetized condition. **D**, **E**, and **F** plot the gastric emptying against the amplitude, frequency, and velocity of gastric peristalsis across animals and conditions (red: awake; blue: anesthetized), respectively. The dashed lines indicate the linear fit of their relationships with the dashed line indicating the 95% confidence intervals. The box plots show the median, 25th, and 75th percentiles. The plus signs indicate outliers. Solid circles indicate individual samples. Asterisks indicate statistical significance (paired t-test, * p < 0.05, ** p < 0.01, *** p < 0.001, n.s.: not significant).

### Differences in intestinal filling, transit, and motility

We also observed profound differences between the awake and anesthetized conditions in the small intestine. Although our imaging and analysis focused more on the stomach, sufficient food or nutritional content also entered the small intestine, including duodenum, jejunum, and ileum, allowing us to assess intestinal filling, transit, and motility. From MRI images, the intestinal compartments initially appeared as elongated and tubular segments, which were similar for both awake and anesthetized conditions (Figure 3A & C, left). About halfway into the experiment, for awake rats, the intestinal segments expanded further to receive more content from the stomach, and transited the food downstream (Figure 3A, right), resulting in a slight and non-significant (p = 0.18) increase in intestinal volume by 4.8±3.4%. In the anesthetized condition, the intestinal segments received less food and significantly decreased their collective volume by 10.2±2.2% (p<0.001). Anesthesia also compromised intestinal motility. Although intestinal contractions were observable in both awake and anesthetized conditions, the contractions were more occlusive, driving faster transit in the awake condition than in the anesthetized condition. See Supplementary videos 1, 2, and 3 for a demonstrative example.

## Discussion

In this study, we established the feasibility of performing GI MRI on awake rats by acclimating them to the MRI environment through repeated behavioral training over one week. For the first time, we report MRI-based assessments of gastric emptying, antral motility, and intestinal filling in awake rats following a naturalistic meal, and compare these assessments with those obtained from anesthetized rats given low-dose isoflurane and dexmedetomidine. Our findings indicate that awake rats exhibited faster gastric emptying, stronger antral contractions, more rapid peristaltic propagation, and more effective intestinal filling and transit, whereas the anesthesia compromised, but not entirely diminished, these motor functions. Our protocols and findings help lay the foundation for using MRI in translational imaging in gastroenterology, mitigating anesthesia as a major confounding factor when translating findings from animals to humans ^28^.

The one-week acclimation is central to the success of GI MRI in awake rats. The acclimation involves repeatedly handling and restraining rats over multiple days, facilitating their adaptation ^55,56^ and avoiding GI complications due to acute stress ^57–61^. GI MRI based on gradient echo induces acoustic noise, posing additional challenges. The acoustic noise during GI MRI is typically lower than that during brain fMRI, which often uses echo planar imaging with higher gradients and faster switching. Since awake rodents can successfully acclimate to fMRI scans despite the louder noise ^29–32^, it is expected that they can tolerate the noisy environment during GI MRI, following similar acclimation. Nevertheless, it would be ideal to eliminate the acoustic noise, for example, by using silent MRI based on the zero echo time (ZTE) looping star pulse sequence ^62,63^.

Our physiological evidence suggests that rats can stay awake during GI MRI without experiencing excessive stress. First, the respiratory and heart rates remained stable during imaging for nearly one hour and were in the range of unstressed awake rats reported by others ^29,48,49^. Second, the body movements were intermittent, occurring approximately 21% of the time during GI MRI, and tended to decrease over time. Third, the respiration rate significantly dropped from 99.8±1.9 cpm at acclimation day 1 to 91.0±1.9 cpm during imaging (p < 0.01).

Last, the corticosterone levels immediately before and after the first awake MRI session were similar to the baseline measures during acclimation (before rats were placed in the mock MRI scanner on days 2, 4, and 6, Supplementary Figure 2). These observations suggested that the animals were not under significant stress during imaging. However, our sample size is limited and future studies are desirable to further evaluate the stress level during imaging.

Our findings call for caution when studying GI motility in anesthetized rats. In particular, a high dose (2.5%) of isoflurane could significantly stall the stomach and disrupt peristalsis, potentially through multiple sites or mechanisms of action. Isoflurane may disrupt electromechanical coupling by smooth muscle cells ^66,67^, interfere with gastric pace-making activity through interstitial cells of Cajal ^68^, or suppress neural activity in peripheral or central nervous systems ^69,70^. It is notable that the gastric emptying rate in awake rats was twice that in anesthetized rats (23.7±1.4% compared to 11.2±1.7% emptied at 48 min), which correlates well with the greater amplitudes of peristaltic waves (reduced gastric circumference by 40.7±2.3% in awake compared to 21.7±1.4% in anesthetized rats, Figure 6D). The frequencies of the peristaltic waves were not reduced, being 5.1±0.1cpm in awake, compared to 5.4±0.1cpm in anesthetized rats. Slow waves, on which the peristaltic waves depend, are generated in ICC and communicated electrically to the smooth muscle ^71^. The results suggest that anesthesia did not compromise the ability of ICC to generate slow waves, because the peristaltic frequency was not reduced. On the other hand, the results suggest that the contractile capacity of the gastric muscle was reduced by anesthesia. This could be a direct effect on the muscle, or could be an action on excitatory neurons that determine the amplitudes of peristaltic waves during gastric emptying ^72^. Inhibition of neurons on excitatory neuromuscular transmission is suggested by previous studies ^66,67^. Our observation that isoflurane reduced HRV but not heart rate also suggests an action on autonomic neurons. It is thus important to consider the type and dosage of anesthesia in studies of GI physiology, with the ideal approach being the use of awake animals.

Our protocol, established in male SD rats, may be generalized to female, other strains, species, or disease models in future studies. Of particular interest is to perform GI MRI with awake mice. For mice, there exist a variety of transgenic models and genetic tools ^73,74^. Combining them with awake GI MRI will open a new window of opportunity for interrogating molecular, cellular, circuit, and network mechanisms underlying GI physiology and pathophysiology.

Our study had limitations. First, we initially anesthetized rats and then allowed recovery by removing isoflurane. Such a recovery from anesthesia may differ from naturally awake conditions. It is possible that there are residual effects of the initial anesthesia ^17^. Second, the awake acclimation involved food/drink rewards. This reward may possibly precondition the animals and affect GI physiology. Third, longer scans in the awake condition are particularly useful for adapting this technique to animal models of gastroparesis, which involves further delayed gastric emptying ^64,65^. It is desirable to acclimate and scan awake rats for more than one hour, e.g., four hours. Lastly, our current analysis focused more on the stomach. Future work may further assess other parts of the GI tract for a more complete understanding of GI functions under awake vs. anesthetized conditions.

## Acknowledgments

We thank Angela Yee for her efforts and assistance in animal behavior training and MRI experiments.

## References

1. Coulie B, Tack J, Sifrim D, Andrioli A, Janssens J. Role of nitric oxide in fasting gastric fundus tone and in 5-HT1 receptor-mediated relaxation of gastric fundus. Am J Physiol-Gastrointest Liver Physiol. 1999;276(2):G373–G377. doi:10.1152/ajpgi.1999.276.2.G373

2. Monroe MJ, Hornby PJ, Partosoedarso ER. Central vagal stimulation evokes gastric volume changes in mice: a novel technique using a miniaturized barostat. Neurogastroenterol Motil. 2004;16(1):5–11. doi:10.1046/j.1365-2982.2003.00464.x

3. Aghababaie Z, Cheng LK, Paskaranandavadivel N, et al. Targeted ablation of gastric pacemaker sites to modulate patterns of bioelectrical slow wave activation and propagation in an anesthetized pig model. Am J Physiol-Gastrointest Liver Physiol. 2022;322(4):G431–G445. doi:10.1152/ajpgi.00332.2021

4. Alighaleh S, Cheng LK, Angeli-Gordon TR, O’Grady G, Paskaranandavadivel N. Optimization of Gastric Pacing Parameters Using High-Resolution Mapping. IEEE Trans Biomed Eng. 2023;70(10):2964–2971. doi:10.1109/TBME.2023.3272521

5. Lu KH, Cao J, Oleson ST, Powley TL, Liu Z. Contrast-Enhanced Magnetic Resonance Imaging of Gastric Emptying and Motility in Rats. IEEE Trans Biomed Eng. 2017;64(11):2546–2554. doi:10.1109/TBME.2017.2737559

6. Lu K -H., Cao J, Oleson S, et al. Vagus nerve stimulation promotes gastric emptying by increasing pyloric opening measured with magnetic resonance imaging. Neurogastroenterol Motil. 2018;30(10):e13380. doi:10.1111/nmo.13380

7. Wang X, Cao J, Han K, et al. Diffeomorphic Surface Modeling for MRI-Based Characterization of Gastric Anatomy and Motility. IEEE Trans Biomed Eng. 2023;70(7):2046–2057. doi:10.1109/TBME.2023.3234509

8. Anderson DL, Bartholomeusz FD, Kirkwood ID, et al. Liquid Gastric Emptying in the Pig: Effect of Concentration of Inhaled Isoflurane. J Nucl Med. 2002;43(7):968–971. Accessed June 21, 2024. https://jnm.snmjournals.org/content/43/7/968

9. Gómez-Lado N, Seoane-Viaño I, Matiz S, et al. Gastrointestinal Tracking and Gastric Emptying of Coated Capsules in Rats with or without Sedation Using CT imaging. Pharmaceutics. 2020;12(1):81. doi:10.3390/pharmaceutics12010081

10. Asai T, Mapleson WW, Power I. Differential effects of clonidine and dexmedetomidine on gastric emptying and gastrointestinal transit in the rat. Br J Anaesth. 1997;78(3):301–307. doi:10.1093/bja/78.3.301

11. Inada T, Asai T, Yamada M, Shingu K. Propofol and Midazolam Inhibit Gastric Emptying and Gastrointestinal Transit in Mice. Anesth Analg. 2004;99(4):1102. doi:10.1213/01.ANE.0000130852.53082.D5

12. Tomaselli L, Sciullo M, Fulton S, et al. Isoflurane anesthesia suppresses gastric myoelectric power in the ferret. Neurogastroenterol Motil. 2024;36(3):e14749. doi:10.1111/nmo.14749

13. Tanila H, Kauppila T, Taira T. Inhibition of intestinal motility and reversal of postlaparotomy ileus by selective *α*2-adrenergic drugs in the rat. Gastroenterology. 1993;104(3):819–824. doi:10.1016/0016-5085(93)91018-D

14. Ailiani AC, Neuberger T, Brasseur JG, et al. Quantifying the effects of inactin vs Isoflurane anesthesia on gastrointestinal motility in rats using dynamic magnetic resonance imaging and spatio-temporal maps. Neurogastroenterol Motil. 2014;26(10):1477–1486. doi:10.1111/nmo.12410

15. Desmet M, Vander Cruyssen P, Pottel H, et al. The influence of propofol and sevoflurane on intestinal motility during laparoscopic surgery. Acta Anaesthesiol Scand. 2016;60(3):335–342. doi:10.1111/aas.12675

16. Schnoor J, Unger JK, Kochs B, Silny J, Rossaint R. Effects of a single dose of ketamine on duodenal motility activity in pigs. Can Vet J. 2005;46(2):147–152. Accessed June 21, 2024. https://www.ncbi.nlm.nih.gov/pmc/articles/PMC1082863/

17. Torjman MC, Joseph JI, Munsick C, Morishita M, Grunwald Z. Effects of isoflurane on gastrointestinal motility after brief exposure in rats. Int J Pharm. 2005;294(1-2):65–71. doi:10.1016/j.ijpharm.2004.12.028

18. Chang H, Li S, Li Y, et al. Effect of sedation with dexmedetomidine or propofol on gastrointestinal motility in lipopolysaccharide-induced endotoxemic mice. BMC Anesthesiol. 2020;20(1):227. doi:10.1186/s12871-020-01146-z

19. Li Y, Wang Y, Chang H, et al. Inhibitory Effects of Dexmedetomidine and Propofol on Gastrointestinal Tract Motility Involving Impaired Enteric Glia Ca2+ Response in Mice. Neurochem Res. 2021;46(6):1410–1422. doi:10.1007/s11064-021-03280-7

20. Botman J, Hontoir F, Gustin P, et al. Postanaesthetic effects of ketamine-midazolam and ketamine-medetomidine on gastrointestinal transit time in rabbits anaesthetised with isoflurane. Vet Rec. 2020;186(8):249. doi:10.1136/vr.105491

21. Keller J, Bassotti G, Clarke J, et al. Advances in the diagnosis and classification of gastric and intestinal motility disorders. Nat Rev Gastroenterol Hepatol. 2018;15(5):291–308. doi:10.1038/nrgastro.2018.7

22. Marciani L. Assessment of gastrointestinal motor functions by MRI: a comprehensive review. Neurogastroenterol Motil. 2011;23(5):399–407. doi:10.1111/j.1365-2982.2011.01670.x

23. Chavero-Pieres M, Viola MF, Appeltans I, et al. Magnetic resonance imaging as a non-invasive tool to assess gastric emptying in mice. Neurogastroenterol Motil. 2023;35(2):e14490. doi:10.1111/nmo.14490

24. Marciani L, Gowland PA, Spiller RC, et al. Effect of meal viscosity and nutrients on satiety, intragastric dilution, and emptying assessed by MRI. Am J Physiol-Gastrointest Liver Physiol. 2001;280(6):G1227–G1233. doi:10.1152/ajpgi.2001.280.6.G1227

25. Kwiatek MA, Steingoetter A, Pal A, et al. Quantification of distal antral contractile motility in healthy human stomach with magnetic resonance imaging. J Magn Reson Imaging JMRI. 2006;24(5):1101–1109. doi:10.1002/jmri.20738

26. Sclocco R, Nguyen C, Staley R, et al. Non-uniform gastric wall kinematics revealed by 4D Cine magnetic resonance imaging in humans. Neurogastroenterol Motil. 2021;33(8):e14146. doi:10.1111/nmo.14146

27. Lu KH, Liu Z, Jaffey D, et al. Automatic assessment of human gastric motility and emptying from dynamic 3D magnetic resonance imaging. Neurogastroenterol Motil. 2022;34(1):e14239. doi:10.1111/nmo.14239

28. Wang X, Alkaabi F, Choi M, et al. Surface Mapping of Gastric Motor Functions Using MRI: A Comparative Study between Humans and Rats. Am J Physiol-Gastrointest Liver Physiol. Published online June 25, 2024. doi:10.1152/ajpgi.00045.2024

29. King JA, Garelick TS, Brevard ME, et al. Procedure for minimizing stress for fMRI studies in conscious rats. J Neurosci Methods. 2005;148(2):154–160. doi:10.1016/j.jneumeth.2005.04.011

30. Ferris CF, Smerkers B, Kulkarni P, et al. Functional magnetic resonance imaging in awake animals. 2011;22(6):665–674. doi:10.1515/RNS.2011.050

31. Zhang N, Rane P, Huang W, et al. Mapping resting-state brain networks in conscious animals. J Neurosci Methods. 2010;189(2):186–196. doi:10.1016/j.jneumeth.2010.04.001

32. Liang Z, King J, Zhang N. Uncovering Intrinsic Connectional Architecture of Functional Networks in Awake Rat Brain. J Neurosci. 2011;31(10):3776–3783. doi:10.1523/JNEUROSCI.4557-10.2011

33. Desai M, Kahn I, Knoblich U, et al. Mapping brain networks in awake mice using combined optical neural control and fMRI. J Neurophysiol. 2011;105(3):1393–1405. doi:10.1152/jn.00828.2010

34. Becerra L, Pendse G, Chang PC, Bishop J, Borsook D. Robust Reproducible Resting State Networks in the Awake Rodent Brain. PLOS ONE. 2011;6(10):e25701. doi:10.1371/journal.pone.0025701

35. Berns GS, Brooks A, Spivak M. Replicability and Heterogeneity of Awake Unrestrained Canine fMRI Responses. PLOS ONE. 2013;8(12):e81698. doi:10.1371/journal.pone.0081698

36. Belcher AM, Yen CC, Stepp H, et al. Large-Scale Brain Networks in the Awake, Truly Resting Marmoset Monkey. J Neurosci. 2013;33(42):16796–16804. doi:10.1523/JNEUROSCI.3146-13.2013

37. Liu JV, Hirano Y, Nascimento GC, Stefanovic B, Leopold DA, Silva AC. fMRI in the awake marmoset: Somatosensory-evoked responses, functional connectivity, and comparison with propofol anesthesia. NeuroImage. 2013;78:186–195. doi:10.1016/j.neuroimage.2013.03.038

38. Harris AP, Lennen RJ, Marshall I, et al. Imaging learned fear circuitry in awake mice using fMRI. Eur J Neurosci. 2015;42(5):2125–2134. doi:10.1111/ejn.12939

39. Hung CC, Yen CC, Ciuchta JL, et al. Functional MRI of visual responses in the awake, behaving marmoset. NeuroImage. 2015;120:1–11. doi:10.1016/j.neuroimage.2015.06.090

40. Schroeder MP, Weiss C, Procissi D, Disterhoft JF, Wang L. Intrinsic connectivity of neural networks in the awake rabbit. NeuroImage. 2016;129:260–267. doi:10.1016/j.neuroimage.2016.01.010

41. Andics A, Gábor A, Gácsi M, Faragó T, Szabó D, Miklósi Á. Neural mechanisms for lexical processing in dogs. Science. 2016;353(6303):1030-1032. doi:10.1126/science.aaf3777

42. Gue M, Fioramonti J, Frexinos J, Alvinerie M, Bueno L. Influence of acoustic stress by noise on gastrointestinal motility in dogs. Dig Dis Sci. 1987;32(12):1411–1417. doi:10.1007/BF01296668

43. Zhang L, Gong JT, Zhang HQ, et al. Melatonin Attenuates Noise Stress-induced Gastrointestinal Motility Disorder and Gastric Stress Ulcer: Role of Gastrointestinal Hormones and Oxidative Stress in Rats. J Neurogastroenterol Motil. 2015;21(2):189–199. doi:10.5056/jnm14119

44. Cao J, Lu KH, Powley TL, Liu Z. Vagal nerve stimulation triggers widespread responses and alters large-scale functional connectivity in the rat brain. PloS One. 2017;12(12):e0189518. doi:10.1371/journal.pone.0189518

45. Cao J, Lu KH, Oleson ST, et al. Gastric stimulation drives fast BOLD responses of neural origin. NeuroImage. 2019;197:200–211. doi:10.1016/j.neuroimage.2019.04.064

46. Cao J, Wang X, Chen J, Zhang N, Liu Z. The vagus nerve mediates the stomach-brain coherence in rats. NeuroImage. 2022;263:119628. doi:10.1016/j.neuroimage.2022.119628

47. Shaffer F, Ginsberg JP. An Overview of Heart Rate Variability Metrics and Norms. Front Public Health. 2017;5:258. doi:10.3389/fpubh.2017.00258

48. Švorc P, Grešová S, Švorc P. Heart rate variability in male rats. Physiol Rep. 2023;11(18):e15827. doi:10.14814/phy2.15827

49. Azar T, Sharp J, Lawson D. Heart Rates of Male and Female Sprague–Dawley and Spontaneously Hypertensive Rats Housed Singly or in Groups. J Am Assoc Lab Anim Sci JAALAS. 2011;50(2):175–184. Accessed June 21, 2024. https://www.ncbi.nlm.nih.gov/pmc/articles/PMC3061417/

50. Svorc P, Svorc P, Gresova S. Sex differences, chronobiology and general anaesthesia in activities of the autonomic nervous system in rats. Exp Physiol. 2023;108(6):810–817. doi:10.1113/EP091143

51. Donoho DL. De-noising by soft-thresholding. IEEE Trans Inf Theory. 1995;41(3):613–627. doi:10.1109/18.382009

52. Thirion JP. Image matching as a diffusion process: an analogy with Maxwell’s demons. Med Image Anal. 1998;2(3):243–260. doi:10.1016/s1361-8415(98)80022-4

53. Vercauteren T, Pennec X, Perchant A, Ayache N. Diffeomorphic demons: efficient non-parametric image registration. NeuroImage. 2009;45(1 Suppl):S61-72. doi:10.1016/j.neuroimage.2008.10.040

54. Indireshkumar K, Brasseur JG, Faas H, et al. Relative contributions of “pressure pump” and “peristaltic pump” to gastric emptying. Am J Physiol-Gastrointest Liver Physiol. 2000;278(4):G604–G616. doi:10.1152/ajpgi.2000.278.4.G604

55. Zheng J, Dobner A, Babygirija R, Ludwig K, Takahashi T. Effects of repeated restraint stress on gastric motility in rats. Am J Physiol-Regul Integr Comp Physiol. 2009;296(5):R1358–R1365. doi:10.1152/ajpregu.90928.2008

56. Zheng J, Babygirija R, Bülbül M, Cerjak D, Ludwig K, Takahashi T. Hypothalamic oxytocin mediates adaptation mechanism against chronic stress in rats. Am J Physiol - Gastrointest Liver Physiol. 2010;299(4):G946–G953. doi:10.1152/ajpgi.00483.2009

57. Enck P, Holtmann G. Stress and gastrointestinal motility in animals: a review of the literature. Neurogastroenterol Motil. 1992;4(2):83–90. doi:10.1111/j.1365-2982.1992.tb00084.x

58. Jiang Y, Greenwood-Van Meerveld B, Johnson AC, Travagli RA. Role of estrogen and stress on the brain-gut axis. Am J Physiol - Gastrointest Liver Physiol. 2019;317(2):G203–G209. doi:10.1152/ajpgi.00144.2019

59. Babygirija R, Zheng J, Bülbül M, Cerjak D, Ludwig K, Takahashi T. Sustained Delayed Gastric Emptying During Repeated Restraint Stress in Oxytocin Knockout Mice. J Neuroendocrinol. 2010;22(11):1181–1186. doi:10.1111/j.1365-2826.2010.02069.x

60. Lewis MW, Hermann GE, Rogers RC, Travagli RA. In vitro and in vivo analysis of the Effects of corticotropin releasing factor on rat dorsal vagal complex. J Physiol. 2002;543(1):135–146. doi:10.1113/jphysiol.2002.019281

61. Sinen O, Akçalı İ, Akkan SS, Bülbül M. The role of hypothalamic Orexin-A in stress-induced gastric dysmotility: An agonistic interplay with corticotropin releasing factor. Neurogastroenterol Motil. 2024;36(1):e14719. doi:10.1111/nmo.14719

62. Wiesinger F, Menini A, Solana AB. Looping Star. Magn Reson Med. 2019;81(1):57-68. doi:10.1002/mrm.27440

63. Xiang H, Fessler JA, Noll DC. Model-based reconstruction for looping-star MRI. Magn Reson Med. 2024;91(5):2104–2113. doi:10.1002/mrm.29927

64. Mard SA, Ahmadi I, Ahangarpour A, Gharib-Naseri MK, Badavi M. Delayed gastric emptying in diabetic rats caused by decreased expression of cystathionine gamma lyase and H2 S synthesis: in vitro and in vivo studies. Neurogastroenterol Motil. 2016;28(11):1677–1689. doi:10.1111/nmo.12867

65. Camilleri M, Chedid V, Ford AC, et al. Gastroparesis. Nat Rev Dis Primer. 2018;4(1):41. doi:10.1038/s41572-018-0038-z

66. Dryn D, Luo J, Melnyk M, Zholos A, Hu H. Inhalation anaesthetic isoflurane inhibits the muscarinic cation current and carbachol-induced gastrointestinal smooth muscle contractions. Eur J Pharmacol. 2018;820:39–44. doi:10.1016/j.ejphar.2017.11.044

67. Zholos AV, Melnyk MI, Dryn DO. Molecular mechanisms of cholinergic neurotransmission in visceral smooth muscles with a focus on receptor-operated TRPC4 channel and impairment of gastrointestinal motility by general anaesthetics and anxiolytics. Neuropharmacology. 2024;242:109776. doi:10.1016/j.neuropharm.2023.109776

68. Aghababaie Z, Wang THH, Nisbet LA, et al. Anaesthesia by intravenous propofol reduces the incidence of intra-operative gastric electrical slow-wave dysrhythmias compared to isoflurane. Sci Rep. 2023;13(1):11824. doi:10.1038/s41598-023-38612-w

69. Zhang X. Effects of Anesthesia on Cerebral Blood Flow and Functional Connectivity of Nonhuman Primates. Vet Sci. 2022;9(10):516. doi:10.3390/vetsci9100516

70. Toader E, Cividjian A, Quintin L. Isoflurane suppresses central cardiac parasympathetic activity in rats: a pilot study. Minerva Anestesiol. 2011;77(2):142–146.

71. Sanders KM, Kito Y, Hwang SJ, Ward SM. Regulation of Gastrointestinal Smooth Muscle Function by Interstitial Cells. Physiol Bethesda Md. 2016;31(5):316–326. doi:10.1152/physiol.00006.2016

72. Wilbur BG, Kelly KA. Effect of proximal gastric, complete gastric, and truncal vagotomy on canine gastric electric activity, motility, and emptying. Ann Surg. 1973;178(3):295–303. Accessed July 10, 2024. https://www.ncbi.nlm.nih.gov/pmc/articles/PMC1355805/

73. Deisseroth K. Optogenetics: 10 years of microbial opsins in neuroscience. Nat Neurosci. 2015;18(9):1213–1225. doi:10.1038/nn.4091

74. Sternson SM, Roth BL. Chemogenetic Tools to Interrogate Brain Functions. Annu Rev Neurosci. 2014;37(Volume 37, 2014):387-407. doi:10.1146/annurev-neuro-071013-014048

