## Supplementary figures and images for "Magnetic Resonance Imaging of Gastric Motility in Conscious Rats"

### supp_figure1_v2.png

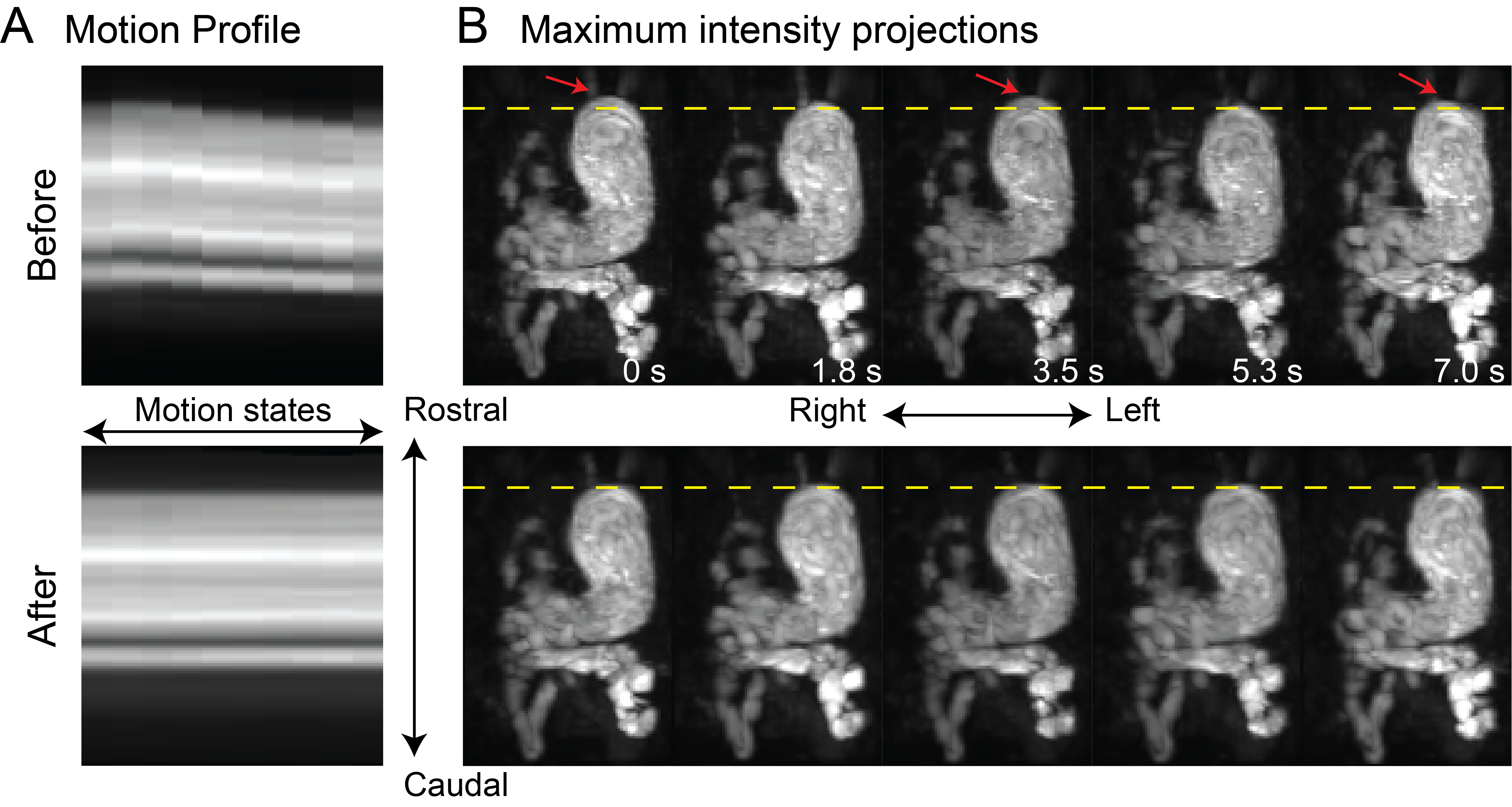

### supp_figure2_v3.png

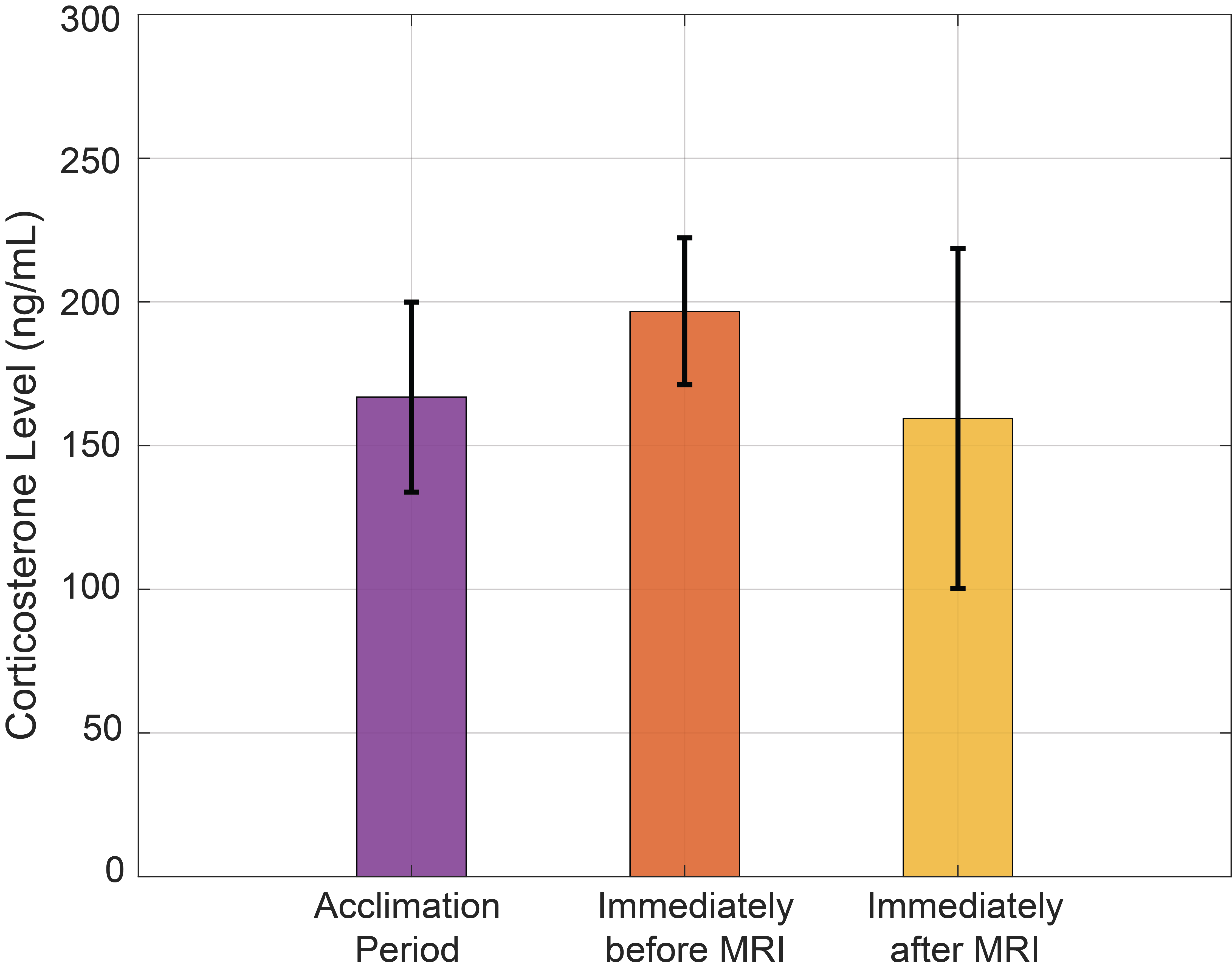

### Supplementary Video 1

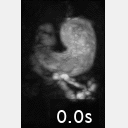

### Supplementary Video 2

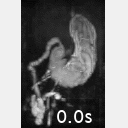

### Supplementary Video 3

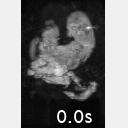
